# A Modified Hodgkin-Huxley model to Show the Effect of Motor Cortex Stimulation on the Trigeminal Neuralgia Network

**DOI:** 10.1101/467100

**Authors:** Mohammadreza Khodashenas, Golnaz Baghdadi, Farzad Towhidkhah

## Abstract

**Background:** Trigeminal neuralgia (TN) is a severe neuropathic pain, which has an electric shock like characteristic. There are some common treatments for this pain such as medicine, microvascular decompression or radio frequency. In this regard, transcranial direct current stimulation (tDCS) is another therapeutic method to reduce the pain, which has been recently attracting the therapists’ attention. The positive effect of tDCS on TN was shown in many previous studies. However, the mechanism of tDCS effect has remained unclear

**Objective:** This study aims to model the neuronal behavior of the main known regions of the brain participating in TN pathways to study the effect of transcranial direct current stimulation

**Method:** The proposed model consists of several blocks (block diagram): 1) trigeminal nerve, 2) trigeminal ganglion, 3) PAG (Periaqueductal gray in the brainstem), 4) thalamus, 5) motor cortex (M1) and 6) somatosensory cortex (S1). Each of these components represented by a modified Hodgkin-Huxley (HH) model (a mathematical model). The modification of the HH model was done based on some neurological facts of pain sodium channels. The input of the model is any stimuli to ‘trigeminal nerve,’ which cause the pain, and the output is the activity of the somatosensory cortex. An external current, which is considered as electrical current, was applied to the motor cortex block of the model

**Result:** The results showed that by decreasing the conductivity of the slow sodium channels (pain channels) and applying tDCS over the M1, the activity of the somatosensory cortex would be reduced. This reduction can cause pain relief

**Conclusion:** The proposed model provided some possible suggestions about the relationship between the effects of tDCS and associated components in TN, and also the relationship between the pain measurement index, somatosensory cortex activity, and the strength of tDCS.

## BACKGROUND

The TN (trigeminal neuralgia) is a rare facial pain disorder that leads to a sudden, short, and severe sense of a pain in the face (1, 2). It is one of the most severe neuropathic pain (3). This pain does not have a regular and normal behavior with a specific pattern. Therefore, the prediction of its occurrence is somehow impossible. It may occur either spontaneously without doing any particular activity or by doing some routine tasks such as chewing, brushing teeth or even shaving, which can trigger the pain attack (1, 2).

Physiological factors (e.g., Superior cerebellar artery compression) and plasticity of the nervous system have roles in TN to play (4). Many diverse regions of the brain such as the thalamus, motor cortex (M1), brainstem, primary somatosensory cortex are included in the TN processing (5, 6), and connections and communications between these regions processing the TN, result in TN network or TN neuromatrix.

Carbamazepine, as one of the TN medicine treatment, can reduce the pain. However, the side effects of this medicine (e.g., drowsiness and confusion) usually results in discontinuation of its usage (1, 7). Surgical interventions such as microvascular decompression or radiotherapies are other options that may be suggested to the patient with TN. Patients often do not tend to have surgery, because it has a high risk of face mutilation (1). Transcranial direct current stimulation (tDCS) is another therapeutic method that is recently used in the field of pain and shows positive effect (1, 8). This method is cheap and non-invasive. No serious side effect has been reported for this method.

The tDCS is a low direct current (usually 1 or 2mA) which is applied to a specific region of the brain using two electrodes, that are placed on the superficial part of the brain. Motor cortex (M1) stimulation is more prevalent than in other regions of the brain. In this regard, M1 stimulation is utilized for pain relief, depression, addiction and so on (9, 10). It was suggested that, by applying tDCS, pain perception is modulated by shifts of the resting membrane potential (1) and consequently results in the modification of the neuronal excitability at the stimulation site (1, 11). Electrical stimulation (e.g., tDCS) of an appropriate area can play a role similar to that of the medial brain in reducing pain (4).

Despite the positive results of the effect of stimulation in pain relief, it is still unknown how tDCS can reduce the symptoms of TN. Modeling the pain pathway can provide a tool to understand some aspects of TN and to investigate the mechanism of tDCS effect. No computational model has been suggested for TN. However, there are some models of pain based on the gate control theory (4) and artificial neural network (12).

In the current study, we simplified a conceptual model of TN pathways that is proposed in the previous study (6). Then we represented this conceptual model by a mathematical formula based on a modified version of Hodgkin-Huxley (HH) equations. By using this model, the possible effects and mechanism of the influence of an external input such as tDCS were investigated. To evaluate the outcomes of our model, as we may not able to understand the meaning of S1 output potential or S1 activity outcome clearly, which is essential for investigating any modification in neuronal behavior in our model, we can change it to a more tangible and practical scale such as visual analogue scale (VAS) to comprehend the intensity of pain and the activity of S1. As a result, interpreting the output of our model, S1 activity is turned to VAS display, a method which is indeed efficient and practical for subjective measuring of pain, including TN. This self-evaluation scale ranges from 0 to 10 as visually described in centimeter units: 0 cm indicates no pain, and 10 cm means the worst pain possible. Participants will be asked to rate their pain during the previous 24 hours to get a baseline pain. This scale has been widely used in studies to evaluate pain as an outcome (13). There are a few types of research which have done some experiments on TN patients by applying tDCS over M1 (1, 8, 14, 15)

In the next section, the details of the proposed conceptual and computational models are presented. The results of the simulation of the proposed model, considering the effect of tDCS, are described in the Results section. In the last part, the discussion and interpretation of the obtained results are provided from different computational and physiological aspects.

## METHOD

In this section, the stages of modeling have been described. At first, a simplified conceptual model of TN pathway has been introduced. This model, which has been explained in (6) in details, consists of some important brain regions involved in TN. Then, each part of the TN pathway has been modeled by a modified version of the HH model. At last step, an external current stimulation has been applied over M1 to show the effect of external stimuli on TN.

### Trigeminal Neuralgia Pathway

Many studies have investigated the brain regions involved in pain processing (1, 8, 16-18). According to the results of these studies, there are a wide variety of brain areas are involved in pain processing that can form a vast network with complex interactions. In our previous work (6), we have described this complicated network as a pain neuromatrix diagram. A simplified version of this neuromatrix is proposed that consists of the leading and substantial blocks of pain network in TN processing system from the initial noxious stimuli of TN to somatosensory cortex (11, 19-23). The simplified pain neuromatrix model has been shown in Fig. 1.

**Figure 1.**
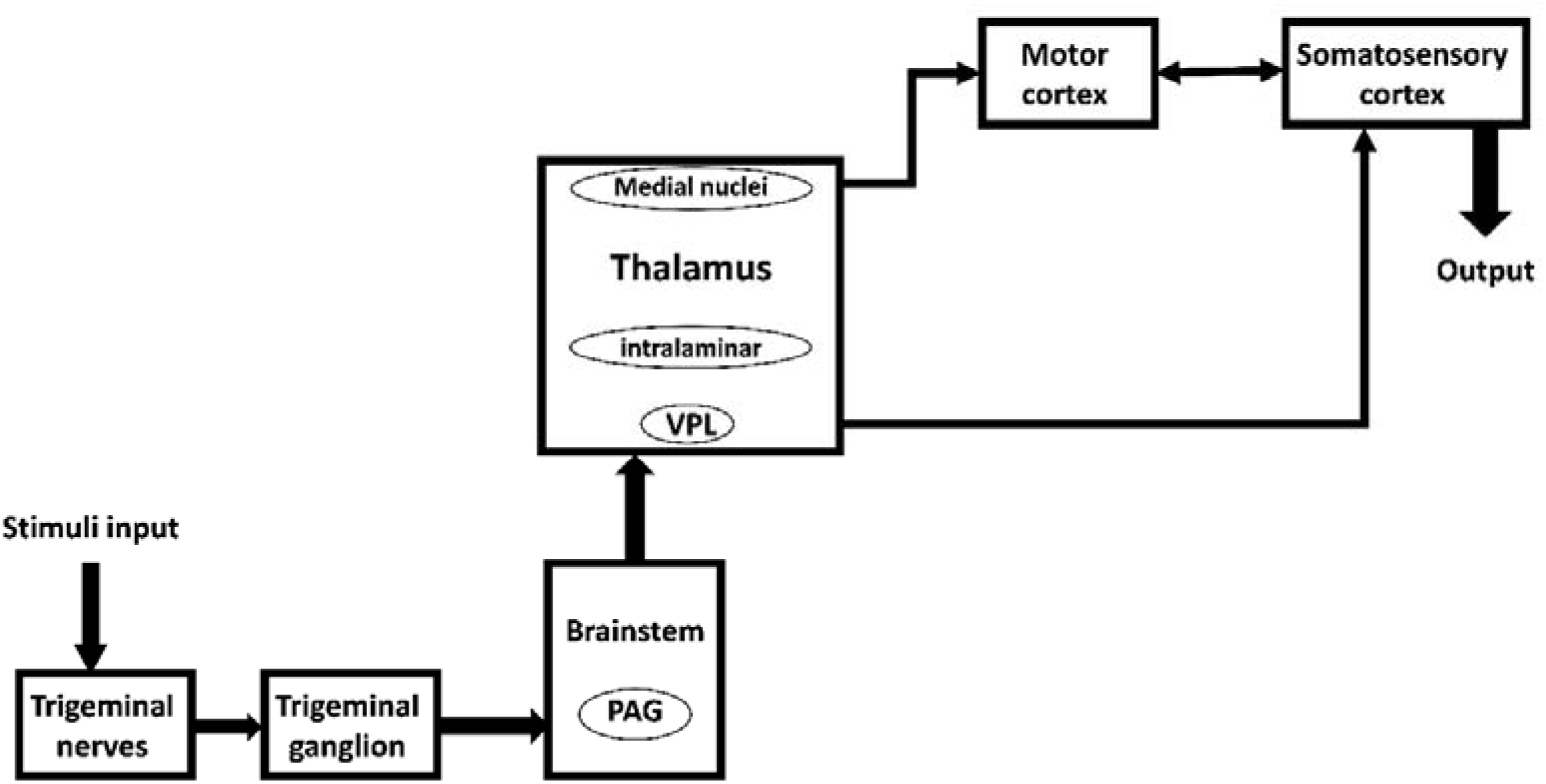
Concise TN pathway block diagram. PAG: periaqueductal gray, VPL: ventral posterolateral nucleus (reprinted from (6)).

As shown in Fig. 1, this model includes the following blocks:

#### Trigeminal ganglion

Trigeminal neuralgia begins from the root of the nerve and trigeminal ganglion (TG) that involved in pain processing pathway. Somas of face neurons are in TG. The signals come from the face, and trigeminal afferents project using the TG, thereby they directly go to the brainstem and then project to the brain (16, 24).

#### Brainstem

After TG, the nociceptive signals reach to different parts of the brainstem (25, 26). Brainstem consists of trigeminal nuclei (16, 18, 27-30), Para brachial (PB) nucleus, and PAG (Periaqueductal gray). Brainstem projects signals to different nuclei of the thalamus (29, 31) especially the VPL (ventral posterolateral nucleus), VPM (ventral posteromedial nucleus) regions (16, 19, 30).

#### Periaqueductal gray (PAG)

Periaqueductal gray is one of the substantial main parts of the pain-mediating process that is in the middle part of the brainstem. It receives signals from thalamus (32), insula, and hypothalamus (31). Periaqueductal gray involves in the secretion of endogenous opioids, such as Encephalin, for relieving pain (12, 19, 26, 28, 31-37).

#### Thalamus

Thalamus is one of the major structures that receives pain signals from diverse pain pathways (18, 19, 25, 26, 28-32, 34-43). Thalamus processes the nociceptive information coming from brainstem (29, 31) especially to VPL and VPM regions (16, 19, 30) and projects them to different parts of the brain such as S2 (secondary somatosensory cortex) (37, 39, 40), primary somatosensory cortex (S1) (19, 30, 31, 37, 39, 40, 42) and PAG (32). It has a reciprocal interaction with some parts of the M1 (35), especially the VL (ventral lateral nucleus) and anterior nuclei (36). In this regard, it has been suggested that the thalamus may have a role in inhibitory pain pathway by applying anodal tDCS over M1, which may result in probable pain relief effect (44).

#### Motor cortex

Although the primary motor cortex (M1) is not considered regularly as one of the pain neuromatrix, it plays a crucial role in modulating the pain in different chronic pain syndromes (25, 28, 35-37, 41, 45). It has some reciprocal connections with S1 (28, 37, 45). It receives direct information from ACC (anterior cingulate cortex) (41) and sends them to the prefrontal cortex (25), brainstem (25, 26) and thalamus (25, 26), and especially VPL (28). Many studies signify the importance and effects of the tDCS over M1 and emphasis on the role of motor cortex stimulation in pain intensity reduction or increase in pain threshold (1, 8, 14, 25, 26, 46-49). Although the mechanism of the effect of the M1 anodal tDCS has remained somehow unclear, such pain relief effects may be because of sub-cortical and thalamo-cortical connections (44).

#### Somatosensory cortex

Primary somatosensory cortex is also one of the main cortical regions in pain or TN neuromatrix (5, 16, 19, 28-31, 35-37, 39-42, 45, 50, 51). Primary somatosensory cortex has some mutual interaction with M1 (28, 37, 45) and S2 (37). Primary somatosensory cortex receives nociceptive information from S2 (41), and thalamus (19, 30, 31, 37, 39, 40, 42).

In the above paragraphs, a brief review of the simplified pain neuromatrix model was provided. More details can be found in (6). In the next section, this model has been formulated by mathematical equations.

### Mathematical Modeling of the Simplified Pain Neuromatrix

Hodgkin–Huxley model, which has been introduced by Hodgkin A. L. and Huxley A. F. (1952), gives the ability to investigate the chemical reactions and activity changes of neuronal response. The equations that describe the HH model, are shown in Eqs. (1)–(10) (52).

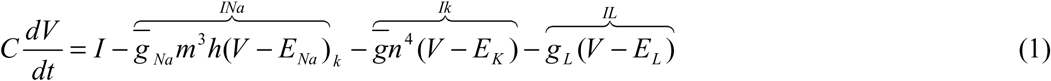

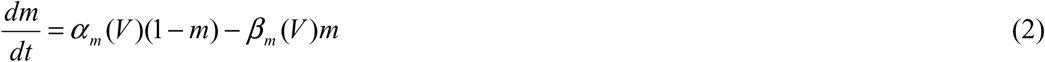

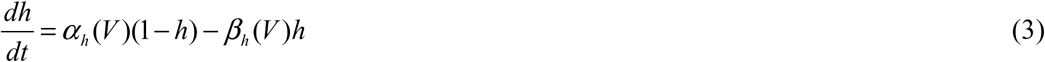

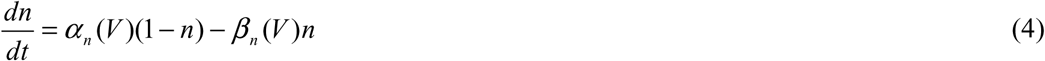

Where

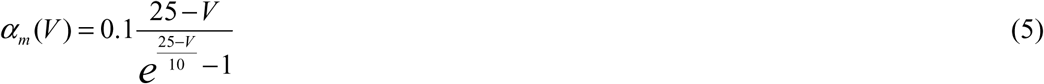

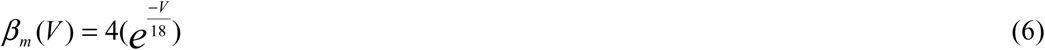

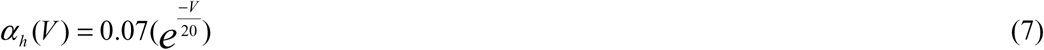

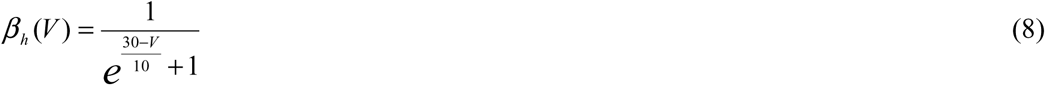

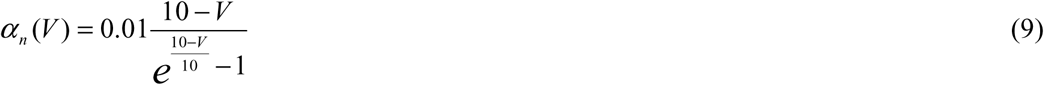

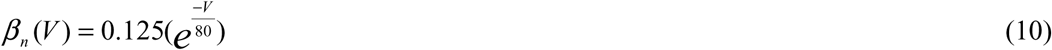

Defenition of variables and parameters invovled in these equations has been provided in Table 1.

**Table 1.** Variables and parameters definitions

| Variable or Parameter Name | Definition |
| --- | --- |
| **E** | Membrane potential |
| **M** | Activation of fast sodium channels |
| **h** | Inactivation of fast sodium channels |
| **m<sub>s</sub>** | Activation of slow sodium channels |
| **h<sub>s</sub>** | Inactivation of slow sodium channels |
| **N** | Activation of potassium channels |
| **C<sub>m</sub>** | Equivalence membrane capacitance |
| **g<sub>Naf</sub>** | Fast sodium channels conductance |
| **g<sub>k</sub>** | Potassium channels conductance |
| **g<sub>l</sub>** | Leak channels conductance |
| **g<sub>NaS</sub>** | Slow sodium channels conductance |
| $E_{Na}$ | Nernst potentials of fast sodium ions |
| $E_k$ | Nernst potentials of potassium ions |
| $E_l$ | Nernst potentials of leakage ions |
| $\alpha_m$ and $\beta_m$ | Transition rates between open and closed states of the activation of sodium channels |
| $\alpha_h$ and $\beta_h$ | Transition rates between open and closed states of the inactivation of sodium channels |
| $\alpha_n$ and $\beta_n$ | Transition rates between open and closed states of the potassium channels |
| $\alpha_{mS}$ and $\beta_{mS}$ | Transition rates between open and closed states of the activation of Slow sodium channels |
| $\alpha_{hS}$ and $\beta_{hS}$ | Transition rates between open and closed states of the inactivation of Slow sodium channels |

The HH model includes three major currents: voltage-gated persistent K^+^ current with four activation gates (resulting in the form of *n*^4^ in the Eq. (1), where *n* is the activation variable for K^+^); voltage-gated transient Na^+^ current with three activation gates and one inactivation gate (indicated by he term *m*^3^*h* in Eq. (1), and leak current, *I*_L_, which is carried by Cl^−^ and other ions (52).

It has been shown that some ion channels, such as the Na_v1.8_ slow sodium channels, have a role in pain pathway and pain intensity modification. In this regard, their synthesis and activity may also cause different neuronal potential and behavoir (53). The HH model has the capability to model and consider the effect of diverse factors influencing on the ion channels. Moreover, the equations presented for HH model can take into account the activity variation of neuronal behaviours. Considering the possible physiological role of activation gating structure of the slow sodium channels Na_*V*_1.8 in impulse coding of nociceptive information (54), and by knowing that the modification of specified slow sodium channels in the membrane of nociceptive neurons is the basis of the pain perception (53), it seems that the HH model is able to be a proper candidate for modeling the pain modulation process. However, it needs some modifications for using in our pain processing study. A voltage-gated slow Na^+^ current needs to be added into the HH equations. Therefore, the modifications have been applied by adding two more equations to the main HH equations (Eqs. (15) and (16)). An extra current for pain intensity plus its corresponding activity fluctuation need to be considered in the HH model. In fact, the added current is the Na_V1.8_ slow sodium channel current specified for pain and pain modulation processing (53). The modified version of HH (MHH) model described by Eqs. (11)- (26).

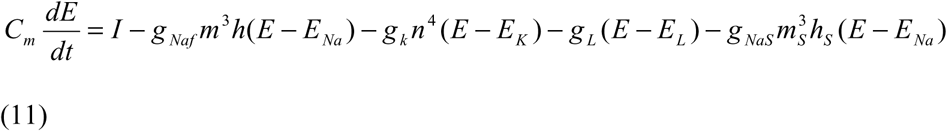

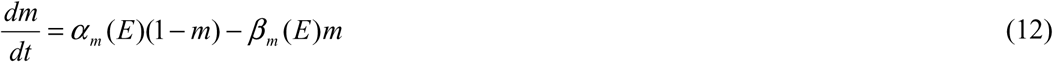

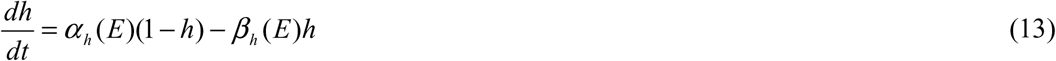

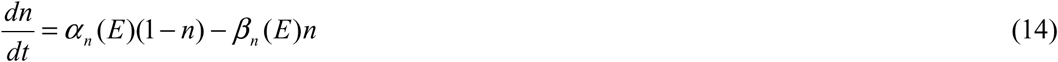

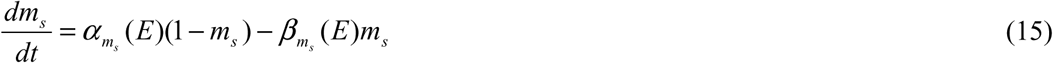

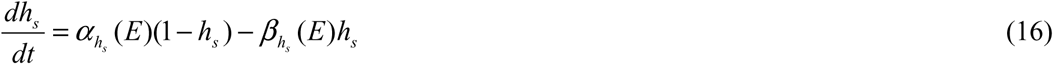

Where

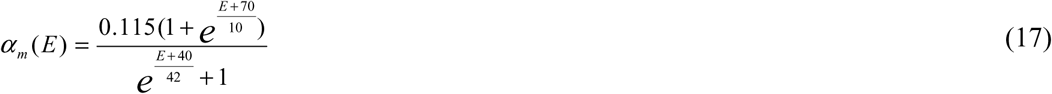

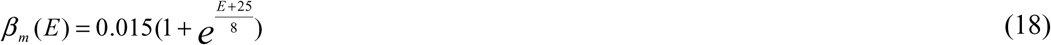

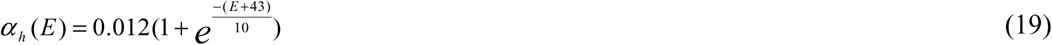

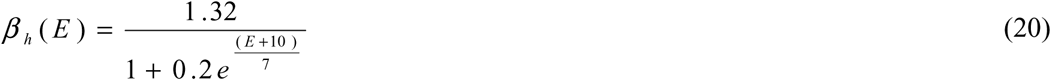

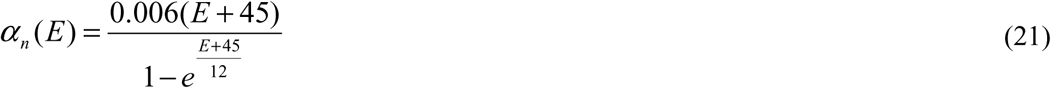

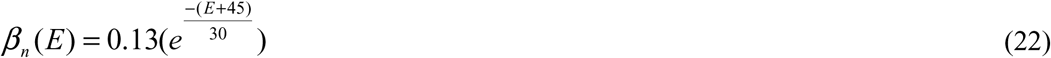

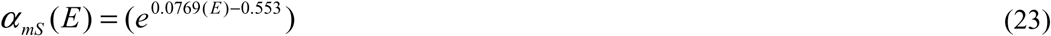

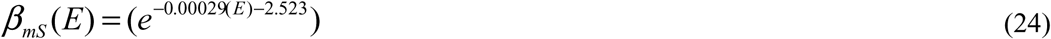

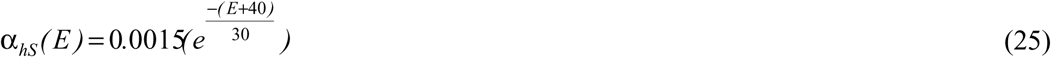

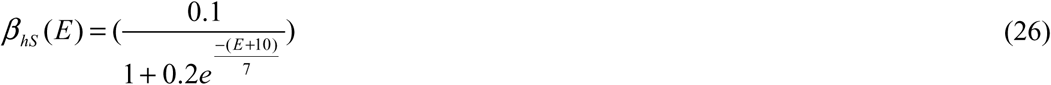

The parameters and variables used in Eqs. (1)-(26) have been introduced in Table 1.

As shown in Fig. 2, to formulate the simplified pain neuromatrix model (Fig. 1), each block was modeled by MHH equations. The input current (*I*) is the noxious stimuli. So, this input was applied to the region where the pain was started (i.e., TG). The characteristics of TN, such as high intensity of pain, shock-like and discontinuity of occurrence, have been represented by the features of the input (*I*).

**Figure 2.**
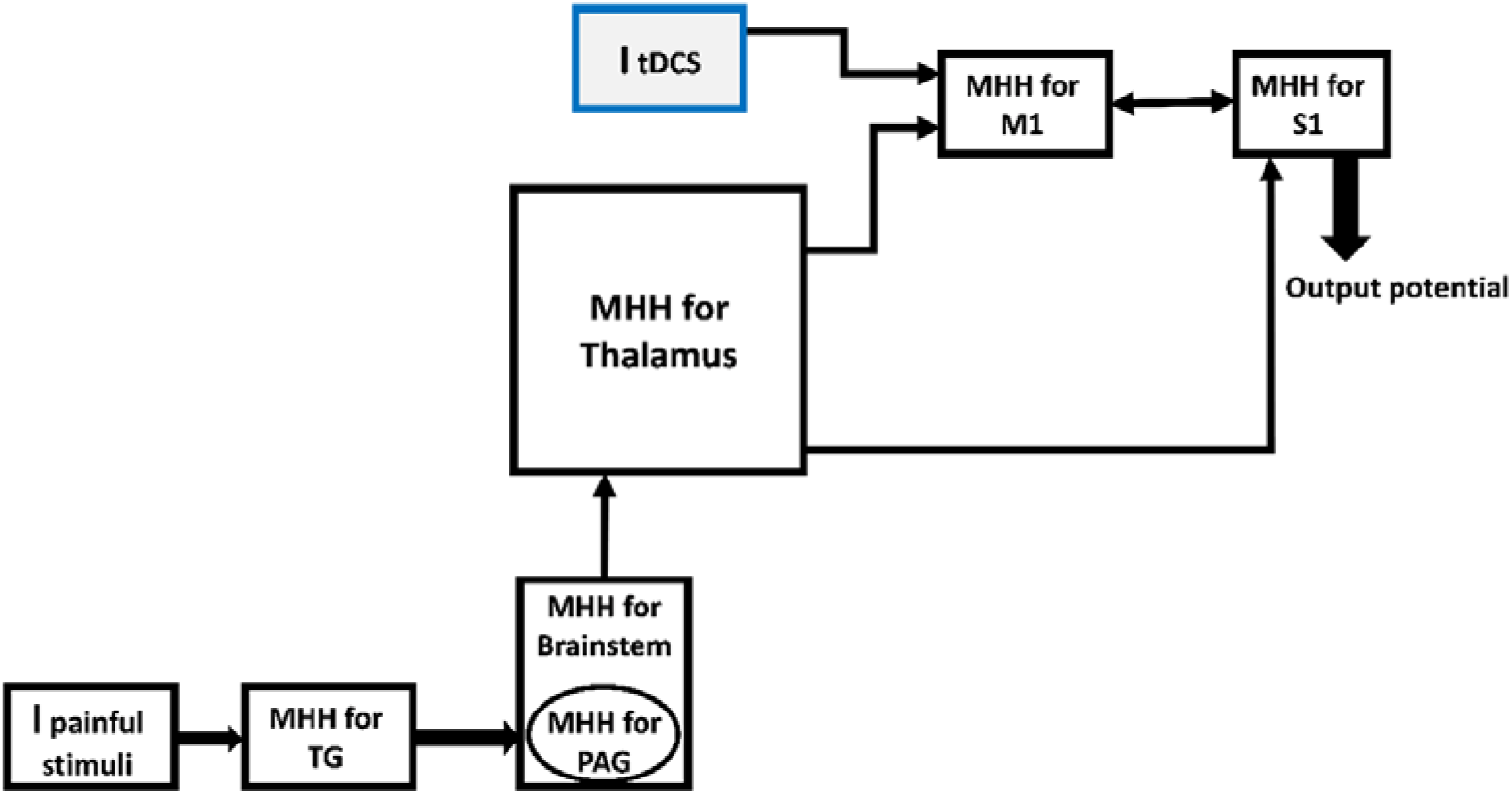
Embedding MHH for each block and inserting ItDCS to M1. MHH: Modified Hodgkin Huxley, TG: Trigeminal Ganglion. PAG: Periaqueductal, M1: Motor Cortex, S1: Somatosensory Cortex.

Each block of the model is differentiated from other ones by considering its specific characteristics such as the membrane capacitances, initial membrane potentials, and conductivities of channels that are proportional to initial potentials of the block. The conductance of each block can be varied from one block to another; however, without losing the whole issue, we considered the same conductance for each block for simplicity. The output *potential* (*E*) of each block is the input of the next block. Therefore, it is required to reform the output potential (*E*) to input current (*I*) of the next block. The mentioned conductance (*G*) is used to transform the output potential (*E*) form of the previous block to the input current form (*I*) of the next block (i.e., *I* = *G* * *V*).

The *capacitance* of blocks is calculated from Eq. (27).

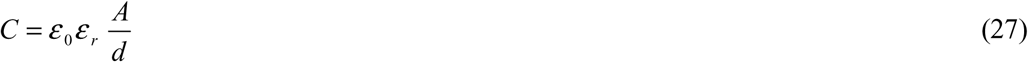

Where, *ε*_0_ = 8.854*10−^12^ and *ε_r_* are the absolute permittivity and relative permittivity of the selected region (i.e., each block) of the brain, respectively. *A* is the area of the membrane cross-section, *d* is a separation between intra and extracellular.

The proposed model has been simulated considering parameters amounts that have been reported in the next part.

Matlab R2013b with SCR:001622 RRID number used as a software tool.

## RESULTS

In the equations which were described in the previous section, the values of the parameters have been selected as indicated in Table 2. The numbers are based on some formula and the amounts reported in previous studies, which have been mentioned beside each amount.

**Table 2.**
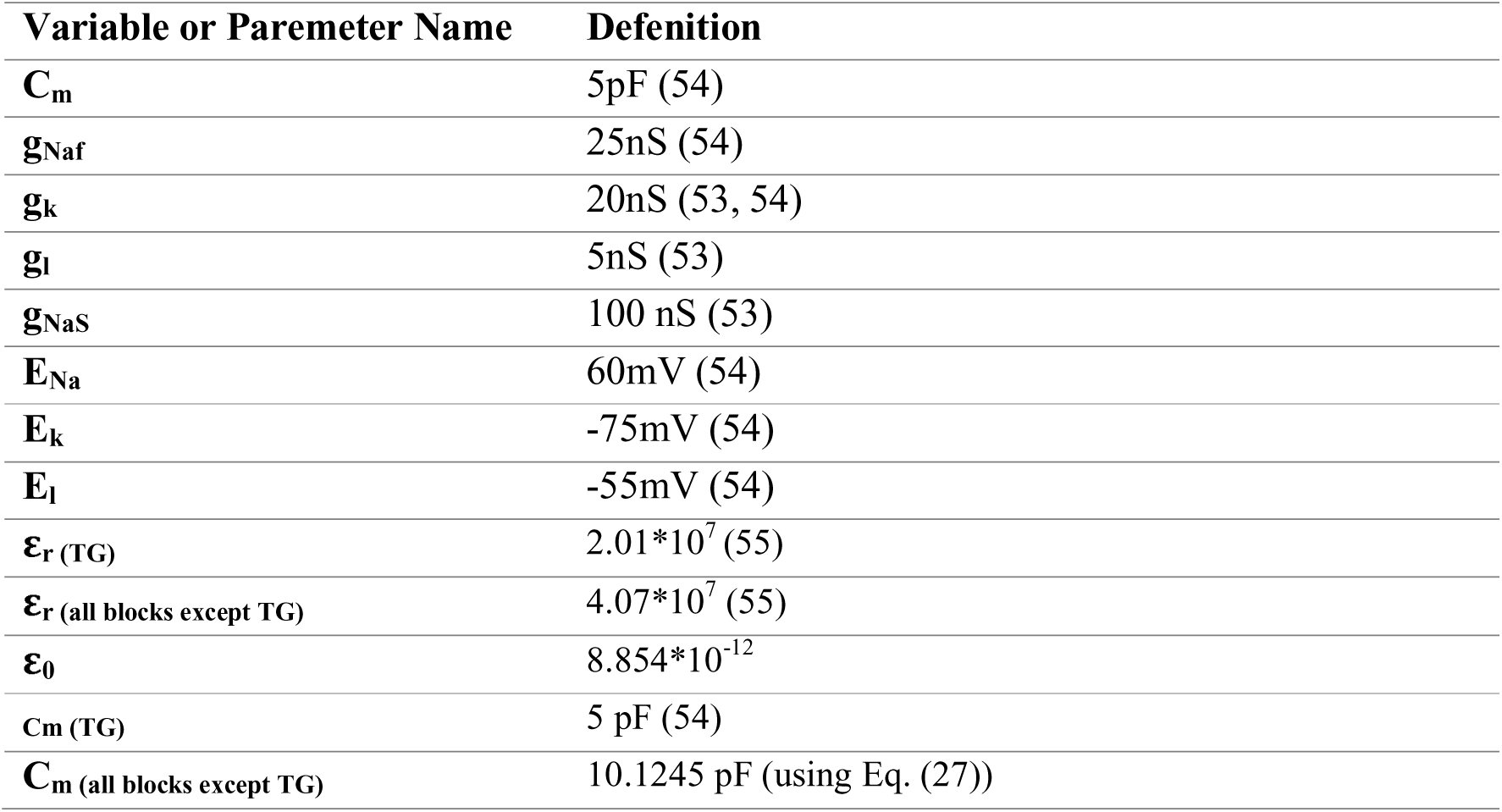
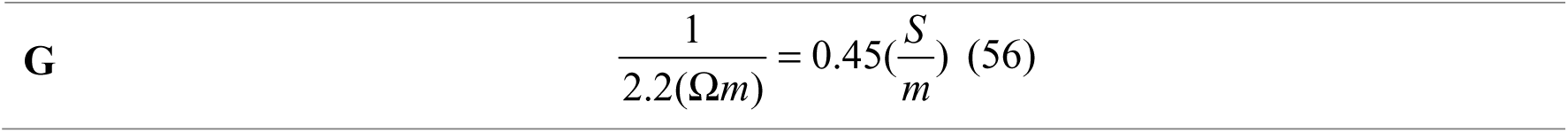
Specific values used for simulation

As shown in Fig. 2, the input stimulus is applied to the TG block. As a result, the simulation indicates that the amplitude of the input affects the output behavior of this block that is shown in Fig. 3.

**Figure 3.**
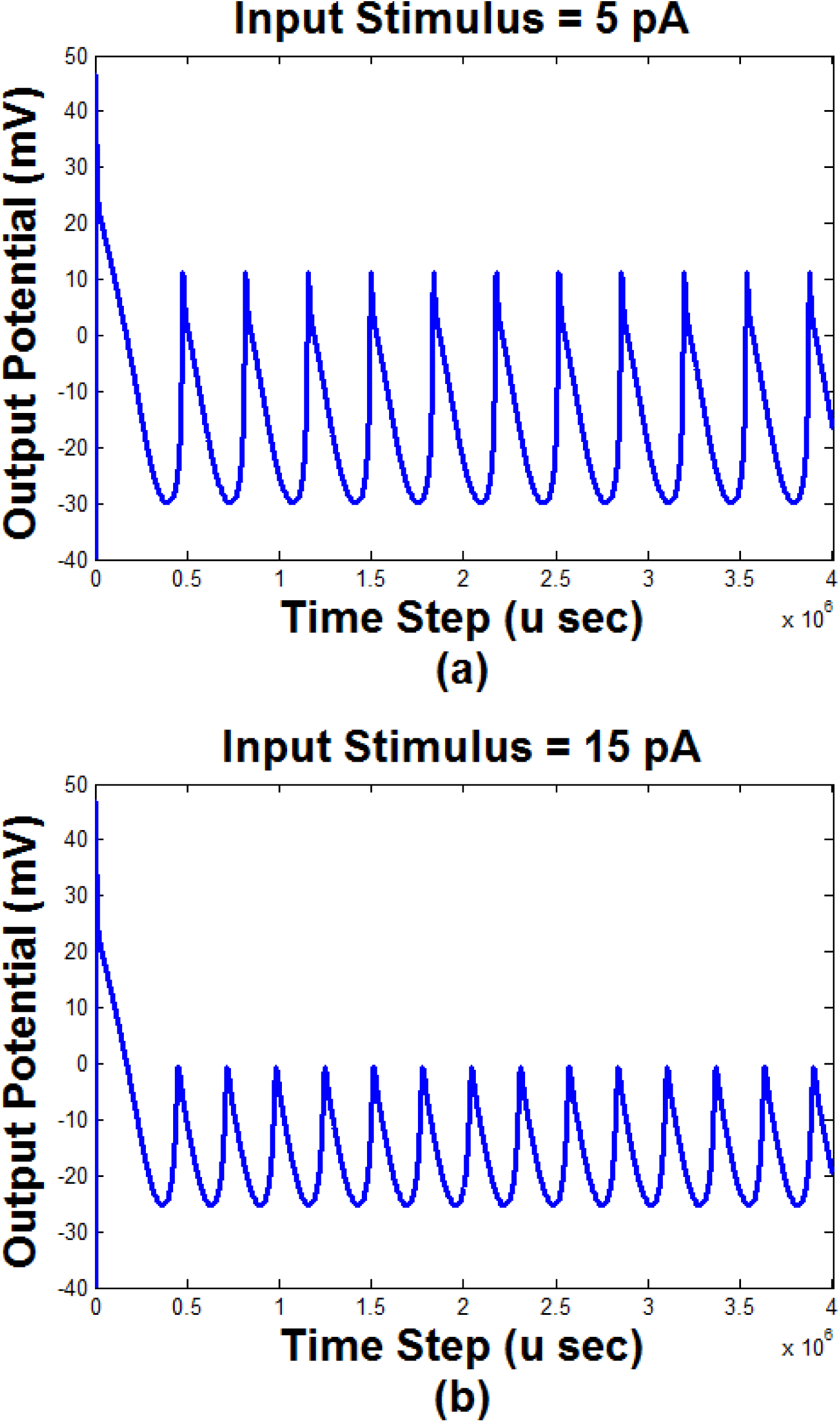
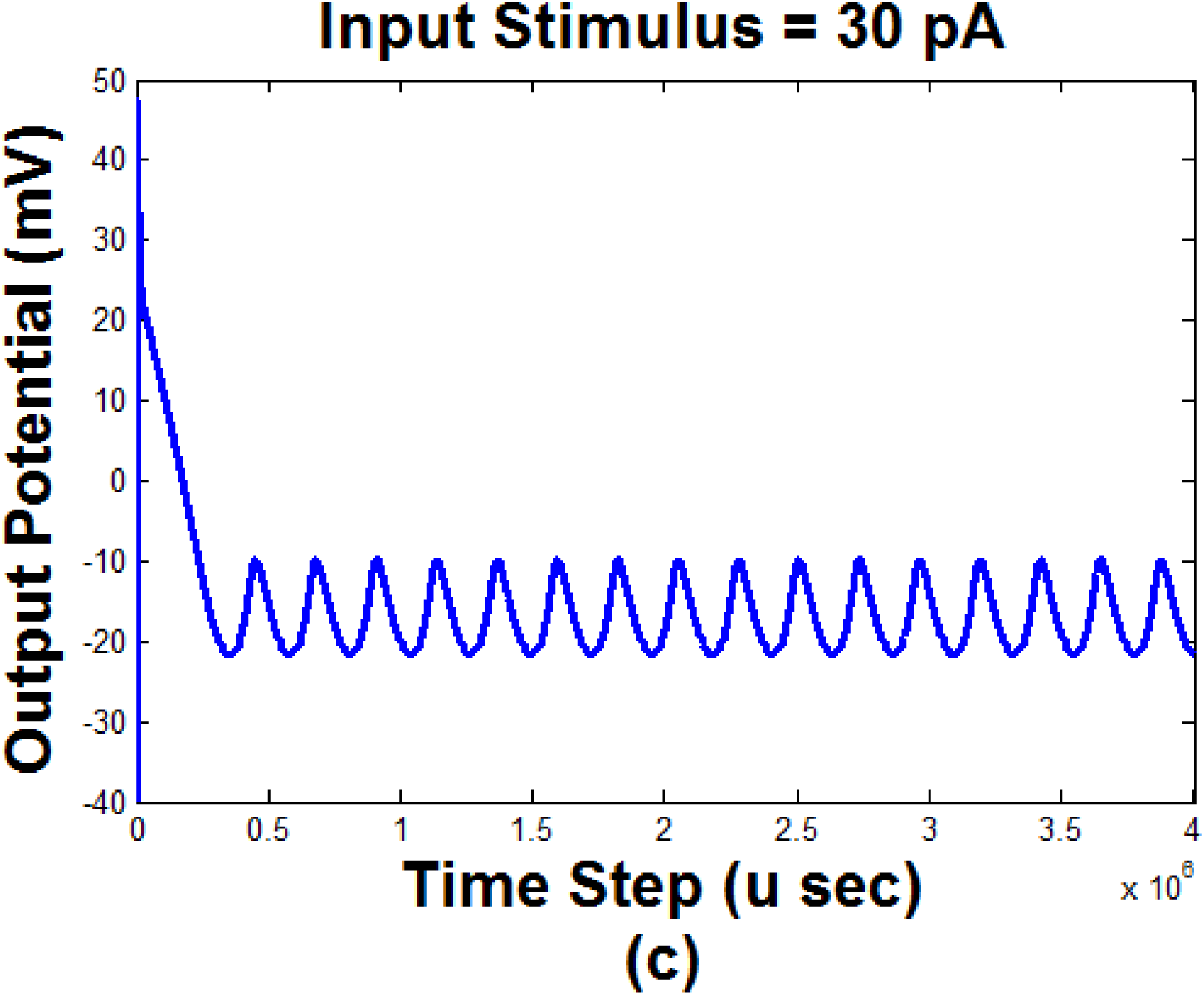
The response of the first block of the model (TG) to the input stimulus with different amplitudes ((a) input stimulus=5pA; (b) input stimulus=15pA; (c) input stimulus=30pA).

Figure 3 demonstrates that applying a different amount of input stimulus changes the behavior of the output of the first block. According to Fig. 3, the peak to peak value of the output decreased by increasing the input stimulus. Regardless of the transient part of the output, the increment of the input stimulus led to the increment of the minimum value of the output and decrement of the maximum value. Increasing the input amplitude, had no considerable effect on the transient part. However, it seems that the upward slope of the output changes declined by the increment of the input stimulus. When the input amplitude is 5pA, Fig.3 (a) shows that the output has 11 peaks (cycles) during the time of the stimulation (i.e., length of the horizontal axis). According to Fig. 3 (b), increasing the input to 15pA, the number of peaks increased to 14. The more increment of the input to 30pA led to the changes in the peak’s number from 14 to 16 (Fig. 3 (c)). Therefore, it can be concluded that increasing the input amplitude (i.e., the strength of the pain) affect the activity and the amplitude of the TG output.

According to the outcomes of previous studies, an ordinary pain can be simulated by a constant input current (53, 54). Figure 3 showed the behavior of the TG block to a common pain (i.e., constant current). However, the pain of TN has not a regular shape. It seems that its pattern changes randomly (1, 2, 57). Therefore, to simulate such a pain, we have considered an input current that randomly fluctuated between zero and a positive or negative value. Figure 4 shows the output of TG and S1 (i.e., the output of the model) blocks to this random input with two maximum values.

**Figure 4.**
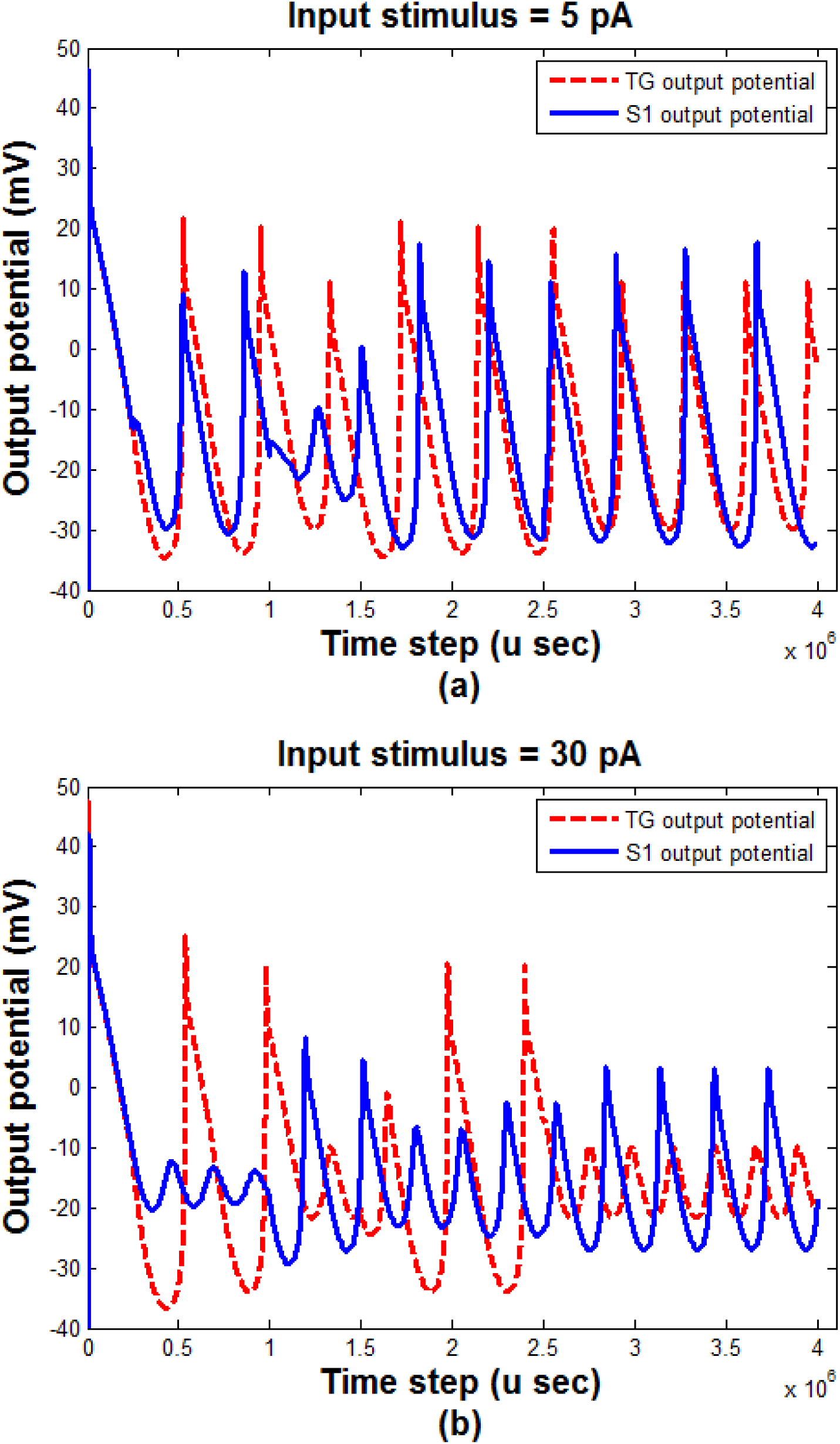
The output of TG and S1 blocks with different input stimulus maximum amplitudes. (a) maximum amplitude=5 pA, (b) maximum amplitude=30 pA. TG: Trigeminal Ganglion, S1: Somatosensory Cortex

As shown in Fig. 4, considering the TN pain the output of the TG block is not as regular as the common pain (Fig. 3). It can be seen that increasing the range of the changes of the input (i.e., TN pain) led to the decrement of the sum squares of the outputs of both TG and S1 blocks. It has also been observed that the phase difference between the output of TG and S1 increases as the range of the input changes is increased.

Figure 5 shows the bifurcation diagram of the extreme values of the S1 output (i.e., model’s output) considering the range of the input changes as the control parameter. The conductivity of the slow sodium channel was g_NaS_=100 nS.

**Figure 5.**
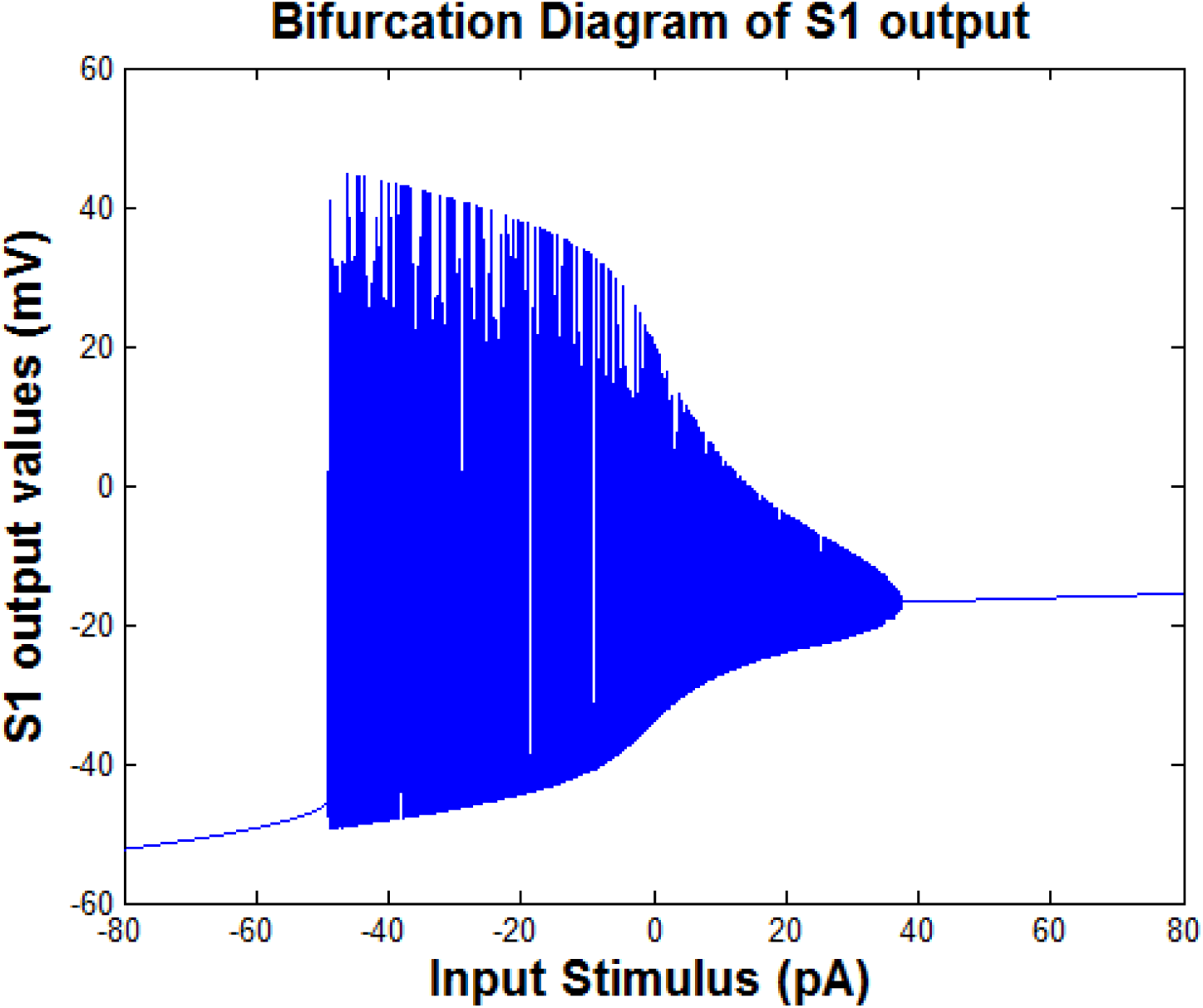
The bifurcation diagram of the values of the S1 output. The range of the input changes is the control parameter (gNaS=100nS)

According to Fig. 5, increasing the range of the input changes upper than about 40pA and decreasing this value lower than about -50pA led to a regular (i.e., harmonic) output. Between these two values (i.e., -50pA and 40pA), the output behavior is bursting and increasing the range of the changes of the input stimulus led to the decrement of the range of the changes of the S1 output. Acceding to the middle part of the diagram (i.e., where the input is between - 50pA to 40pA), it can be seen that the minimum value of the output decreases and the maximum value increases by the increment of the range of input changes.

In addition to the input stimulus, the conductivity of channels can affect the pattern of the output of the model.

Figure 6 shows the effect of the conductivity of slow sodium channels on the output of TG and S1 blocks.

**Figure 6.**
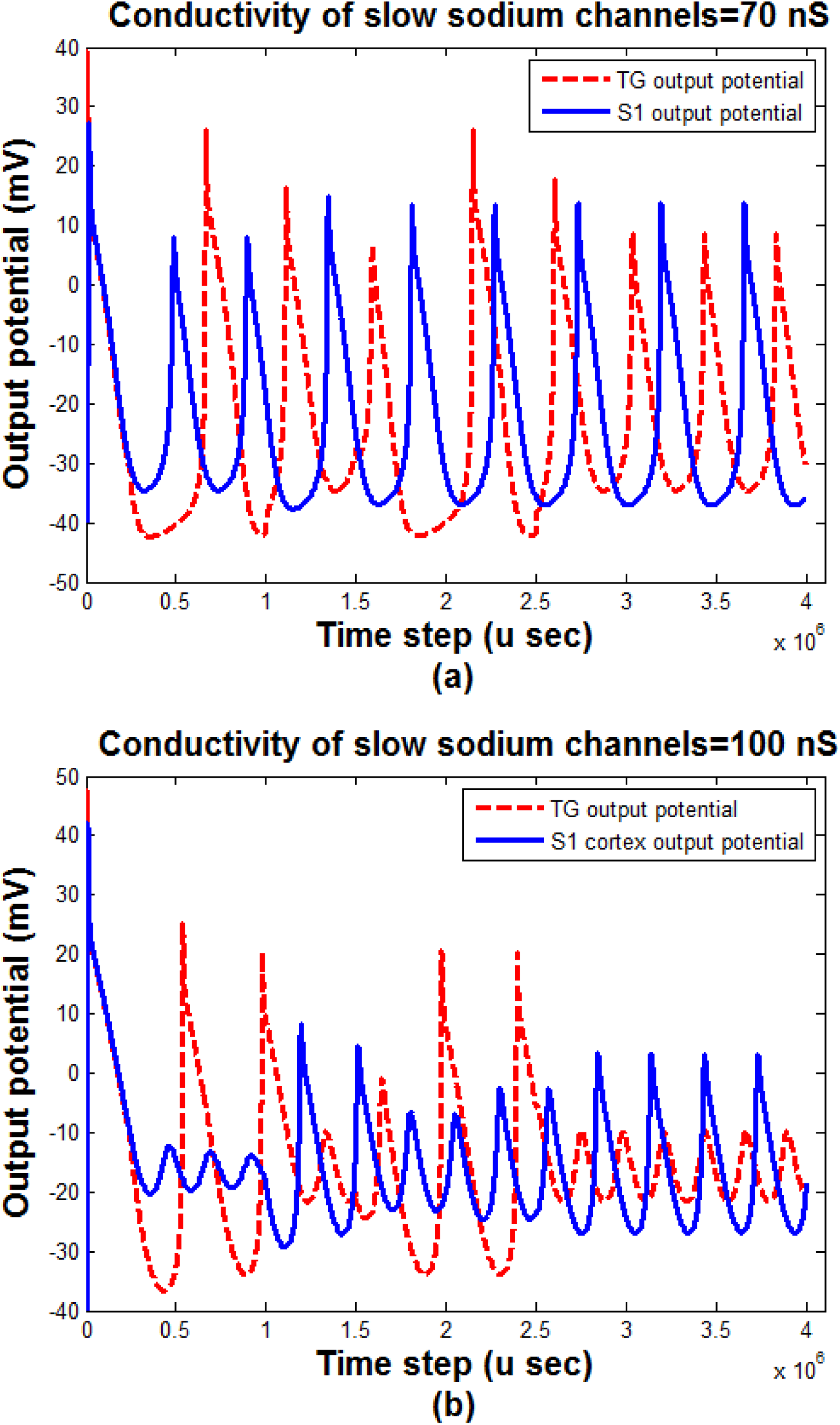
The effect of the conductivity of slow sodium channels on the output pattern of the TG and S1 blocks. (a) Conductivity =70nS, (b) Conductivity = 100nS; input stimulus range=(0,30pA)

According to Fig. 6, the sum squares of the outputs of both TG and S1 blocks is decreased by the increasing of slow sodium channels conductivities named g_NaS_. The phase difference between the outputs of TG and S1 blocks is more obvious in the lower value of the slow sodium channels conductivities than the higher one.

The bifurcation diagram of the values of the S1 output considering the slow sodium channels conductivity as the control parameter is shown in Fig. 7. The input stimulus is considered 30pA.

**Figure 7.**
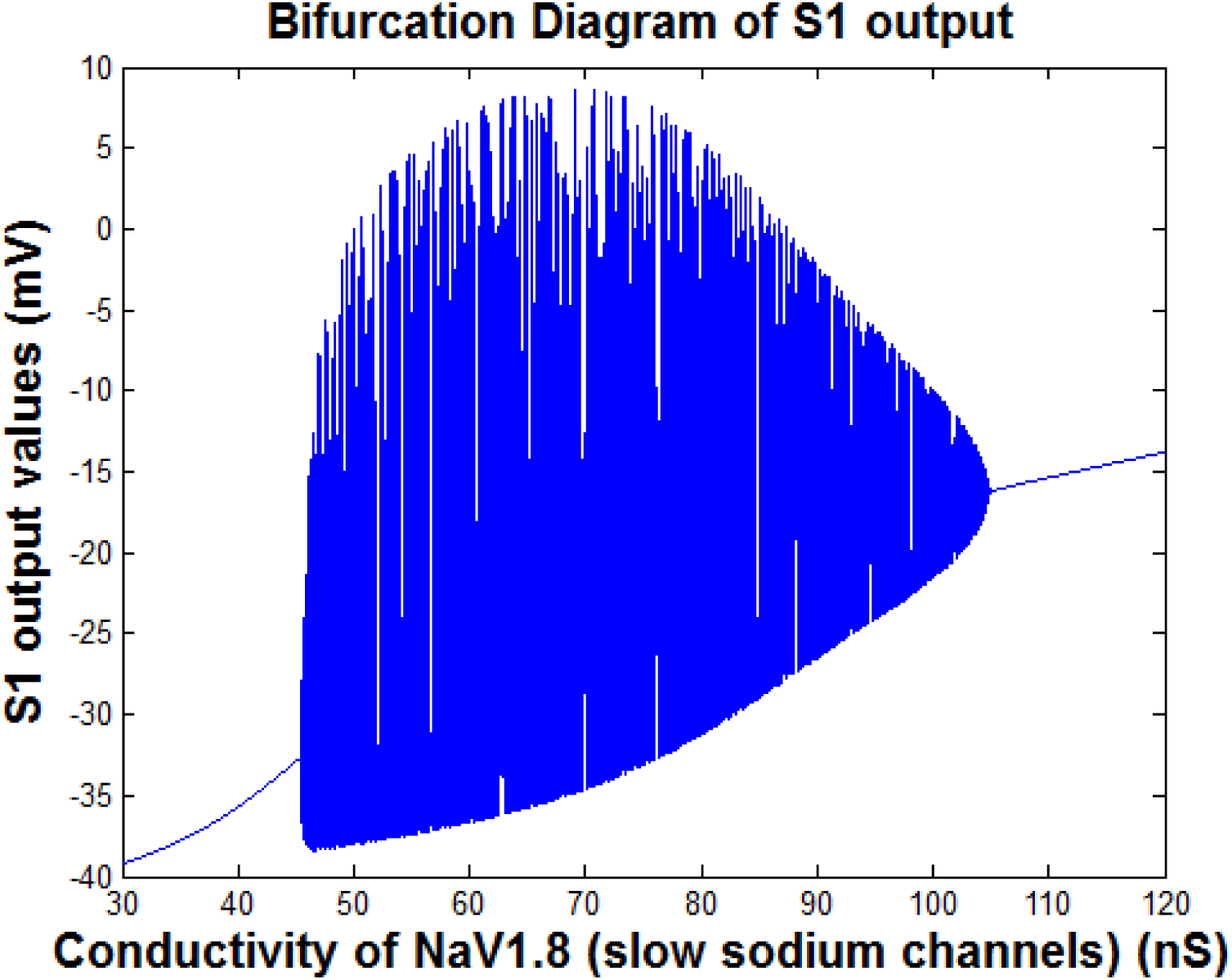
The bifurcation diagram of the extreme values of the S1 output. The slow sodium channels conductivity (gNaS) is the control parameter (input stimulus=30pA)

As shown in Fig. 7, the bifurcation diagram of the S1 output, has three different parts. The first and last parts exhibit a harmonic behavior (with one frequency component) of the output. The amplitude of this harmonic behavior increases by increasing the gNaS. In the middle part of the diagram, the output consists of a variety of frequency component (i.e., bursting behavior) and the minimum value of the output decreases by the increment of the range of input changes. However, the maximum amount of the output has an inverse-U shape. It increases and then decreases by the increment the input range.

The previous simulations were for a model without considering an external current stimulus to M1. By adding another current as an external input current, which can be regarded as a tDCS current (I_tDCS_) to M1 block (seeFig.2), the results of simulations can show the effect of I_tDCS_ on the output behaviors. Figure 8 shows the outputs behavior (i.e., the output of TG and S1 blocks) with and without considering I_tDCS_. In this figure, the input stimulus is 30pA, and the conductivity of pain channels is g_NaS_=100.

**Figure 8.**
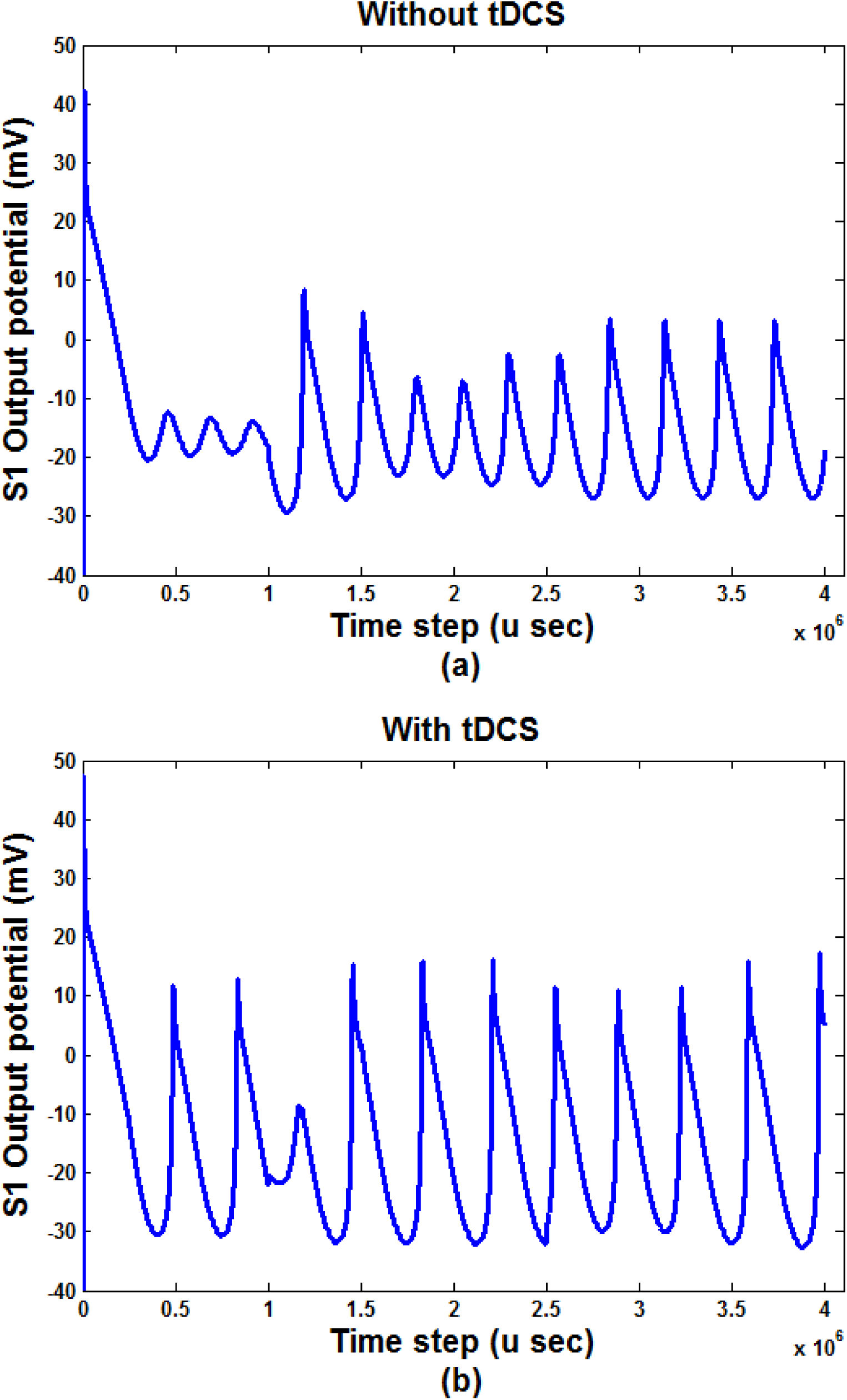
Output behavior of S1 (a) without ItDCS, (b) with ItDCS.

As shown in Fig. 8, the sum squares of the S1 output increased by adding I_tDCS_.

According to the results of the studies (1, 8, 14, 15) done on the effect of tDCS on the pain level (VAS), especially on TN, the points are shown in Fig. 9 are extracted by calculating the average VAS in each current stimulation in all mentioned studies.

**Figure 9.**
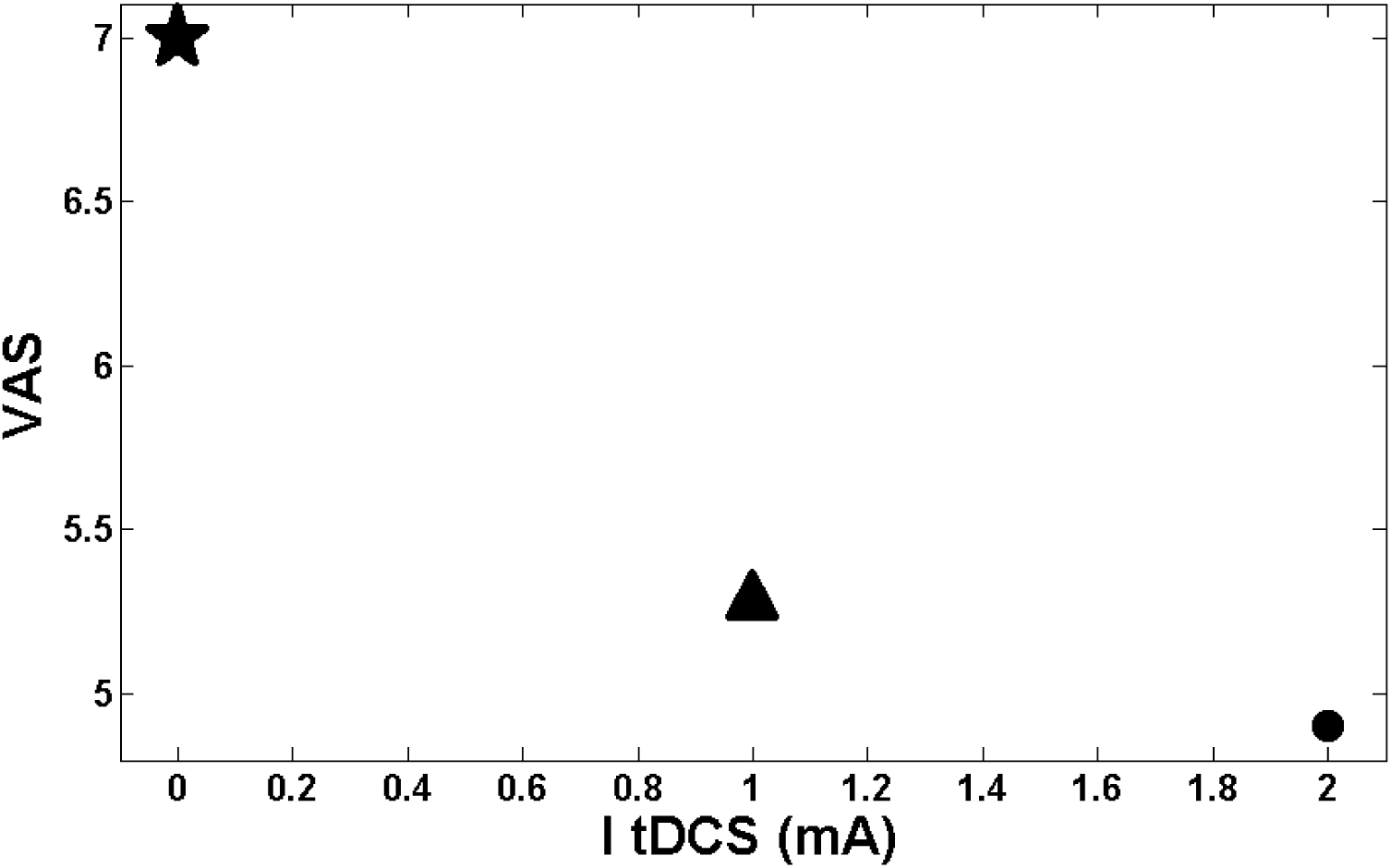
The relationship between ItDCS and the mean value of VAS. The star point shows the mean level of VAS when there is no stimulation obtained from (1, 8, 14). The triangle point shows the mean level of VAS when I_tDCS_= 1 mA obtained from (1, 8, 14); The circular point shows the mean level of VAS when I_tDCS_=2mA obtained from (15);

According to Fig. 9, increasing the level of I_tDCS_ led to the decrement of the pain level (VAS) exponentially.

By using the simulation results of the proposed model, the relationship between the rate of

S1 activity and I_tDCS_ level has been shown in Fig. 10.

**Figure 10.**
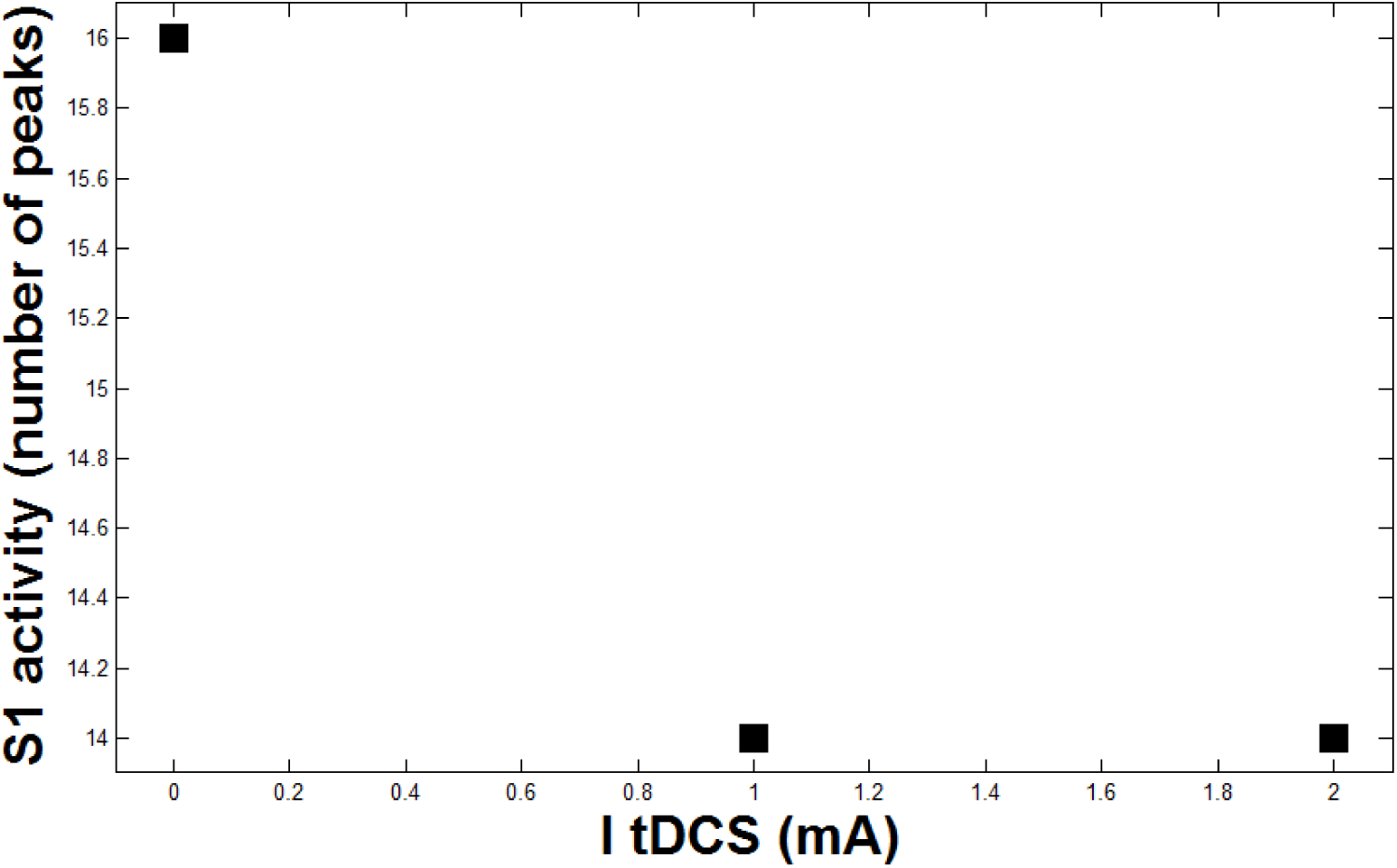
Output of model according to ItDCS

Figure 10 shows an exponential relationship between the S1 activity and the strength of I_tDCS_. As discussed in the introduction part, 1) the pain level is determined by the VAS; 2) electrical stimulation (I_tDCS_) affects the pain level, and 3) the TN as a neuropathic pain changes the pattern of the activities of the neurons (53). Considering these three mentioned points and combining the results presented in Figs. 9 and 10, an exponential relationship between S1 activity and VAS index is obtained (see Fig. 11)

**Figure 11.**
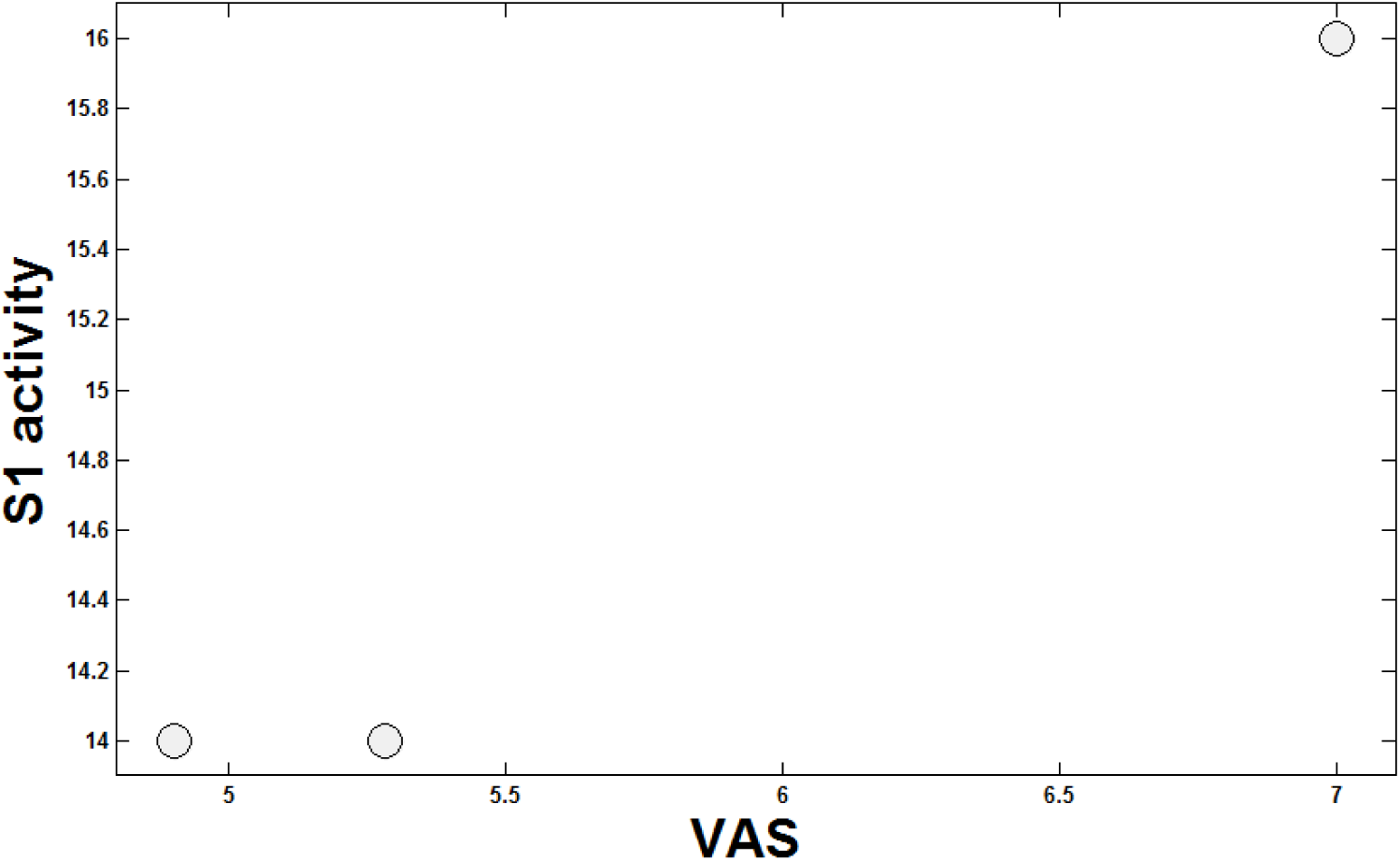
Map of S1 output on VAS

According to Fig. 11, increasing the S1 activity can lead to the increment of the pain level (VAS).

## DISCUSSION

TN is one the most severe pain known among other neuropathic pains but the treatment of it, is still a challenge. Today, different treatment methods such as drugs, microvascular decompression, or surgeries are applied to reduce TN pain. Recently, tDCS has attracted scientific attention as a tool for pain reduction, which can be considered a safe, cheap, and accessible intervention. It has been believed that it may have influences on neurotransmitters (e.g., Glutamate or GABA) and ion concentration in intracellular and extracellular environments. For example, anodal tDCS, which is usually an excitatory stimulation, results in GABA reduction (58). On the other hand, the cathodal stimulation, which is generally an inhibitory stimulation, causes a decrease of Glutamatergic neuronal activity with a highly correlated decrease in GABA (58, 59). Despite the mentioned suggestions, it is still unknown how tDCS affect neural processing. In this study, we aimed to provide some possible ideas about the relationship between the effects of tDCS and associated components in TN, using a computational model. According to the results of this study, as shown in Fig. 6, the improper synthesis of proteins in particular channels (Na_V1.8_) may lead to a decrease in the sensation of neuropathic pain. Besides, the relationship between the activity of these channels and the pain, noxious mediation may cause abnormal sensitivity states like hyperalgesia, which may be treated by diminishing the activity of Na_V1.8_ (53). The catastrophic pain of TN may stem from the activity of these sodium channels in the trigeminal ganglion. Also, pain features affect the response frequency of involved neural systems by influencing neurochemical interactions.

The proposed computational model includes some of the main regions involved in TN. Each of these regions was modeled by a modified version of the HH differential equations, and the results of the simulations demonstrated some effects of the pain stimulus and tDCS on the activity patterns of the model’s components.

According to Fig. 3, increasing the strength of the input stimulus (i.e., pain) led to the decrement of the peak to peak output potential value and the increment of the output frequency (i.e., a number of peaks). That is, as shown in Fig. 3, a further increment of the pain stimulation does not necessarily result in the increase of the S1 activity and consequently pain levels. It seems that there is a nonlinear relationship between the activity of S1 and the strength of the pain stimulus.

The trigeminal ganglion and somatosensory neuronal potential behavior are shown in Fig.4. While the noxious stimuli are increased, the output peak-to-peak activities of both TG and S1 blocks would be increased, as well. The augmentation of the number of peaks and the amplitude of the activities was observed. Therefore, it can be speculated that the more intensifying pain sensation in S1 and TG, the more annoyance and irritation for the patients would occur.

The conductivity of pain channels, g_NaS_, is an influential and substantial parameter of pain propagation in the neuronal pathway(53). The outputs of TG and S1 regions, correspond to the two different pain channels conductivity, are shown in Fig.6. The slow sodium channels, pain channels, synthesis, and activity are reduced by the decrement of Na_v1.8_ channels conductivity. Thereby, the transmission of pain signals may be propagated by decreasing the activity of g_NaS_, which may lead to pain relief. Incidentally, the amplitude variation of outputs (e.g., Fig.6) is because of TN input pattern fluctuation. As we can see in Fig.6 (b), the existence and absence of TN are more obvious than Fig.6 (a) with a lower conductivity of pain channels g_NaS_. In other words, the different peak-to-peak values in different time steps are stemmed from the existence and absence of TN.

In Fig. 8, by applying an external electrical current to the motor cortex block, we see the effect of external current stimulation. In this case, the motor cortex has two inputs. One input comes from the previous block, and another input is the electrical current stimulation over the M1 cortex. We called the second one as I_tDCS_. Somatosensory and motor cortices are connected functionally and structurally (6). Therefore, they can influence each other and cause to a reduction in the sensation of pain. In this regard, the essential role of the motor cortex for alleviating pain will be somehow signified, and such neuronal and functional connections between M1 and other pain-related regions of the brain, especially S1, deemed to be the most crucial part of discussing pain relief by applying external electrical stimulation. In other words, by applying anodal stimulation over M1 area, the sub-cortical, cortico-cortical, thalamo-cortical and connections in the brain may be modulated, which can stem to increase the activity of significant circuits as inhibitory pain pathways, e.g., thalamus and PAG, in the brain, which then causes to pain relief. As shown in Fig. 8, by applying I_tDCS_, the somatosensory cortex (S1) potentials activity is reduced. Therefore, it can be suggested that the pain signals in the region of S1 will be less sensed that is consistent with the reported results of the pain relief by tDCS (8).

Another capability of the model is its potential to map the S1 activities into the VAS value. That is, if we get the output potential activity of S1 from the model, we can approximately estimate the VAS value. Then, we can find the amount of applied stimulation to have a desired amount of VAS.

## CONCLUSION

The developed pain neuromatrix of TN consisted of main regions of the brain that were modeled by an MHH model. By the current version of the model, the possible effect of increasing the pain strength and also the external current stimulus on the TN neuromatrix components were investigated. For future work, other interventions (e.g., transcranial alternating current stimulation (tACS)) to other blocks of the model is suggested. The values of the conductivity and the capacitance could be specified for each block separately in a future study to have a more accurate model.

## LIST OF ABBREVIATIONS

ACC: anterior cingulate cortex
HH: Hodgkin-Huxley
M1: primary motor cortex
MHH: modified Hodgkin-Huxley
PAG: periaqueductal gray
PB: parabrachial nucleus
S1: primary somatosensory cortex
S2: secondary somatosensory cortex
tACS: transcranial alternating current stimulation
tDCS: transcranial direct current stimulation
TG: trigeminal ganglion
TN: trigeminal neuralgia
VAS: visual analogue scale
VPL: ventral posterolateral nucleus
VPM: ventral posteromedial nucleus

## Declarations

### ETHICS APPROVAL AND CONSENT TO PARTICIPATE

Not applicable.

### CONSENT FOR PUBLICATION

Not applicable.

### AVAILABILITY OF DATA AND MATERIAL

Not applicable.

### COMPETING INTERESTS

The authors declare that they have no competing interests.

### FUNDING

This work was partially supported by cognitive science and technologies council (#2572).

### AUTHORS’ CONTRIBUTIONS

MK made substantial contributions to conception and designed the study. MK wrote the first draft of the manuscript and interpreted the findings as a significant contributor. GB has been involved in drafting the manuscript and revising it critically for important intellectual content. MK and GB organized the study. FT was responsible for the supervision of the study. All authors agreed to be accountable for all aspects of the work. All authors read and approved the final manuscript.

## ACKNOWLEDGMENTS

Not applicable.

## REFERENCES

1. Hagenacker T, Bude V, Naegel S, Holle D, Katsarava Z, Diener H-C, et al. Patient-conducted anodal transcranial direct current stimulation of the motor cortex alleviates pain in trigeminal neuralgia. J Headache Pain. 2014;15:78.

2. Maarbjerg S, Gozalov A, Olesen J, Bendtsen L. Trigeminal neuralgia–a prospective systematic study of clinical characteristics in 158 patients. Headache: The Journal of Head and Face Pain. 2014;54(10): 1574-82.

3. Okeson JP. Bell’s orofacial pains: the clinical management of orofacial pain: Quintessence Publishing Company Chicago, III, USA; 2005.

4. Britton N, Skevington SM. A mathematical model of the gate control theory of pain. Journal of theoretical biology. 1989;137(1):91-105.

5. Obermann M, Rodriguez-Raecke R, Naegel S, Holle D, Mueller D, Yoon M-S, et al. Gray matter volume reduction reflects chronic pain in trigeminal neuralgia. Neuroimage. 2013;74:352-8.

6. Khodashenas. M, Towhidkhah. F, Baghdadi. G. A Conceptual Model of Trigeminal Neuralgia Network and tDCS Pain Reduction Effect. Dev Anesthetics Pain Manag DAPM. 2018;1(2):000506.

7. Graham J, Zilkha K. Treatment of trigeminal neuralgia with carbamazepine: a follow-up study. British Medical Journal. 1966;1(5481):210.

8. Obermann Mark, Bude Vera, Holle Dagney, Naegel Steffen, Hagenacker Tim, Diener Hans-Christoph, et al. Anodal Transcranial Direct Current Stimulation Alleviates Pain In Trigeminal Neuralgia. The Journal of Headache and Pain. 2014;15:58.

9. Kuo M-F, Paulus W, Nitsche MA. Therapeutic effects of non-invasive brain stimulation with direct currents (tDCS) in neuropsychiatric diseases. Neuroimage. 2014;85:948-60.

10. Nitsche MA, Cohen LG, Wassermann EM, Priori A, Lang N, Antal A, et al. Transcranial direct current stimulation: state of the art 2008. Brain stimulation. 2008;1(3):206-23.

11. Jaberzadeh S, Bastani A, Zoghi M. Anodal transcranial pulsed current stimulation: A novel technique to enhance corticospinal excitability. Clinical Neurophysiology. 2014;125(2):344-51.

12. Haeri M, Asemani D, Gharibzadeh S. Modeling of pain using artificial neural networks. Journal of Theoretical Biology. 2003;220(3):277-84.

13. Fregni F, Boggio PS, Lima MC, Ferreira MJ, Wagner T, Rigonatti SP, et al. A sham-controlled, phase II trial of transcranial direct current stimulation for the treatment of central pain in traumatic spinal cord injury. Pain. 2006; 122(1-2): 197-209.

14. Antal A, Terney D, Kühnl S, Paulus W. Anodal transcranial direct current stimulation of the motor cortex ameliorates chronic pain and reduces short intracortical inhibition. Journal of pain and symptom management. 2010;39(5):890-903.

15. Donnell A, Nascimento TD, Lawrence M, Gupta V, Zieba T, Truong DQ, et al. High-definition and non-invasive brain modulation of pain and motor dysfunction in chronic TMD. Brain stimulation. 2015;8(6):1085-92.

16. Wang Y, Li D, Bao F, Ma S, Guo C, Jin C, et al. Thalamic metabolic alterations with cognitive dysfunction in idiopathic trigeminal neuralgia: a multivoxel spectroscopy study. Neuroradiology. 2014;56(8):685-93.

17. Hansen N, Obermann M, Poitz F, Holle D, Diener H-C, Antal A, et al. Modulation of human trigeminal and extracranial nociceptive processing by transcranial direct current stimulation of the motor cortex. Cephalalgia. 2011;31(6):661-70.

18. Obermann M, Yoon M, Ese D, Maschke M, Kaube H, Diener H, et al. Impaired trigeminal nociceptive processing in patients with trigeminal neuralgia. Neurology. 2007;69(9):835-41.

19. Ab Aziz CB, Ahmad AH. The role of the thalamus in modulating pain. 2007.

20. Hooks BM, Mao T, Gutnisky DA, Yamawaki N, Svoboda K, Shepherd GM. Organization of cortical and thalamic input to pyramidal neurons in mouse motor cortex. Journal of Neuroscience. 2013;33(2):748-60.

21. Huang T-N, Chuang H-C, Chou W-H, Chen C-Y, Wang H-F, Chou S-J, et al. Tbr1 haploinsuffidency impairs amygdalar axonal projections and results in cognitive abnormality. Nature neuroscience. 2014;17(2):240-7.

22. Oswald MJ, Tantirigama ML, Sonntag I, Hughes SM, Empson RM. Diversity of layer 5 projection neurons in the mouse motor cortex. Frontiers in cellular neuroscience. 2013;7:174.

23. Mercer JG, Moar KM, Findlay PA, Hoggard N, Adam CL. Association of leptin receptor (OBRb), NPY and GLP-1 gene expression in the ovine and murine brainstem. Regulatory peptides. 1998;75:271-8.

24. Hall JE. Guyton and Hall textbook of medical physiology: Elsevier Health Sciences; 2015.

25. Holsheimer J, Nguyen J-P, Lefaucheur J-P, Manola L. Cathodal, anodal or bifocal stimulation of the motor cortex in the management of chronic pain? Operative Neuromodulation: Springer; 2007. p. 57-66.

26. DaSilva AF, Mendonca ME, Zaghi S, Lopes M, DosSantos MF, Spierings EL, et al. tDCS-lnduced Analgesia and Electrical Fields in Pain-Related Neural Networks in Chronic Migraine. Headache. 2012;52(8): 1283-95.

27. Ellrich J. Trigeminal nociceptive reflexes. MOVEMENT DISORDERS-NEW YORK-. 2002;17(2; SUPP):S41-S4.

28. Ebel H, Rust D, Tronnier V, Böker D, Kunze S. Chronic precentral stimulation in trigeminal neuropathic pain. Acta neurochirurgica. 1996;138(11):1300-6.

29. Dubin AE, Patapoutian A. Nociceptors: the sensors of the pain pathway. The Journal of clinical investigation. 2010;120(11):3760-72.

30. DaSilva AF, Becerra L, Makris N, Strassman AM, Gonzalez RG, Geatrakis N, et al. Somatotopic activation in the human trigeminal pain pathway. Journal of Neuroscience. 2002;22(18):8183-92.

31. Prescott SA. Pain Processing Pathway Models. Encyclopedia of Computational Neuroscience. 2015:2181-7.

32. Valet M, Sprenger T, Boecker H, Willoch F, Rummeny E, Conrad B, et al. Distraction modulates connectivity of the cingulo-frontal cortex and the midbrain during pain—an fMRI analysis. Pain. 2004; 109(3):399-408.

33. Kandel ER, Schwartz JH, Jessell TM, Siegelbaum SA, Hudspeth AJ. Principles of neural science: McGraw-hill New York; 2000.

34. Veinante P, Yalcin I, Barrot M. The amygdala between sensation and affect: a role in pain. Journal of molecular psychiatry. 2013;1(1):9.

35. Saavedra LC, Mendonca M, Fregni F. Role of the primary motor cortex in the maintenance and treatment of pain in fibromyalgia. Medical hypotheses. 2014;83(3):332-6.

36. Vaseghi B, Zoghi M, Jaberzadeh S. Does anodal transcranial direct current stimulation modulate sensory perception and pain? A meta-analysis study. Clinical Neurophysiology. 2014; 125(9): 1847-58.

37. Apkarian AV, Bushnell MC, Treede RD, Zubieta JK. Human brain mechanisms of pain perception and regulation in health and disease. European journal of pain. 2005;9(4):463-.

38. Hsieh J-C, Meyerson BA, Ingvara M. PET study on central processing of pain in trigeminal neuropathy. European Journal of Pain. 1999;3(1):51-65.

39. Patrizi F, Freedman SD, Pascual-Leone A, Fregni F. Novel therapeutic approaches to the treatment of chronic abdominal visceral pain. The Scientific World Journal. 2006;6:472-90.

40. Hofbauer RK, Rainville P, Duncan GH, Bushnell MC. Cortical representation of the sensory dimension of pain. Journal of neurophysiology. 2001;86(1):402-11.

41. Bromm B. Brain images of pain. Physiology. 2001;16(5):244-9.

42. Zhu Y, Lu T. A multi-scale view of skin thermal pain: from nociception to pain sensation. Philosophical Transactions of the Royal Society of London A: Mathematical, Physical and Engineering Sciences. 2010;368(1912):521-59.

43. Ogino Y, Nemoto H, Inui K, Saito S, Kakigi R, Goto F. Inner experience of pain: imagination of pain while viewing images showing painful events forms subjective pain representation in human brain. Cerebral Cortex. 2006;17(5):1139-46.

44. Polanía R, Paulus W, Nitsche MA. Modulating cortico-striatal and thalamo-cortical functional connectivity with transcranial direct current stimulation. Human brain mapping. 2012;33(10):2499-508.

45. Vaseghi B, Zoghi M, Jaberzadeh S. How does anodal transcranial direct current stimulation of the pain neuromatrix affect brain excitability and pain perception? A randomised, double-blind, sham-control study. PloS one. 2015;10(3):e0118340.

46. Fregni F, Gimenes R, Valle AC, Ferreira MJ, Rocha RR, Natalle L, et al. A randomized, sham-controlled, proof of principle study of transcranial direct current stimulation for the treatment of pain in fibromyalgia. Arthritis & Rheumatism. 2006;54(12):3988-98.

47. Jaberzadeh S, Vaseghi B, Zoghi M. Cathodal-tDCS induced reduction in excitability of superficial pain neuromatrix cortices is associated with sensory and pain threshold increases. Brain Stimulation. 2015;8(2):337-8.

48. Knotkova H, Soto E, Leuschner Z, Greenberg A, Stock V, Das D, et al. Transcranial direct current stimulation (tDCS) for the treatment of chronic pain. The Journal of Pain. 2013;14(4):S64.

49. Ngernyam N, Jensen MP, Arayawichanon P, Auvichayapat N, Tiamkao S, Janjarasjitt S, et al. The effects of transcranial direct current stimulation in patients with neuropathic pain from spinal cord injury. Clinical Neurophysiology. 2015;126(2):382-90.

50. Coghill RC, McHaffie JG, Yen Y-F. Neural correlates of interindividual differences in the subjective experience of pain. Proceedings of the National Academy of Sciences. 2003;100(14):8538-42.

51. Albanese M-C, Duerden EG, Rainville P, Duncan GH. Memory traces of pain in human cortex. Journal of Neuroscience. 2007;27(17):4612-20.

52. Izhikevich EM. Dynamical systems in neuroscience: MIT press; 2007.

53. Dick OE, Krylov BV. Bifurcation Analysis of Nociceptive Neurons. 2013.

54. Dik O, Shelykh T, Plakhova V, Nozdrachev A, Podzorova S, Krylov B. Application of bifurcation analysis for determining the mechanism of coding of nociceptive signals. Technical Physics. 2015;60(10): 1545-8.

55. Gabriel C. Compilation of the Dielectric Properties of Body Tissues at RF and Microwave Frequencies. DTIC Document, 1996.

56. Malmivuo J, Plonsey R. Bioelectromagnetism: principles and applications of bioelectric and biomagnetic fields: Oxford University Press, USA; 1995.

57. Manzoni G, Torelli P. Epidemiology of typical and atypical craniofacial neuralgias. Neurological Sciences. 2005;26(2):s65-s7.

58. Nitsche MA, Liebetanz D, Schlitterlau A, Henschke U, Fricke K, Frommann K, et al. GABAergic modulation of DC stimulation-induced motor cortex excitability shifts in humans. European Journal of Neuroscience. 2004;19(10):2720-6.

59. Stagg CJ, Best JG, Stephenson MC, O’Shea J, Wylezinska M, Kincses ZT, et al. Polarity-sensitive modulation of cortical neurotransmitters by transcranial stimulation. Journal of Neuroscience. 2009;29(16):5202-6.

